# Dynamical Scaling and Phase Coexistence in Topologically-Constrained DNA Melting

**DOI:** 10.1101/121673

**Authors:** Y. A. G. Fosado, D. Michieletto, D. Marenduzzo

## Abstract

There is a long-standing experimental observation that the melting of topologically constrained DNA, such as circular-closed plasmids, is less abrupt than that of linear molecules. This finding points to an important role of topology in the physics of DNA denaturation, which is however poorly understood. Here, we shed light on this issue by combining large-scale Brownian Dynamics simulations with an analytically solvable phenomenological Landau mean field theory. We find that the competition between melting and supercoiling leads to phase coexistence of denatured and intact phases at the single molecule level. This coexistence occurs in a wide temperature range, thereby accounting for the broadening of the transition. Finally, our simulations show an intriguing topology-dependent scaling law governing the growth of denaturation bubbles in supercoiled plasmids, which can be understood within the proposed mean field theory.

---

One of the most fascinating aspects of DNA is that its biological function is intimately linked to its local topology [1]. For instance, DNA looping [2, 3] and supercoiling [1, 4, 5] are well-known regulators of gene expression, and a variety of proteins, such as Polymerases, Gyrases and Topoisomerases, can affect genomic function by acting on DNA topology [1, 2].

Fundamental biological processes such as DNA transcription and replication are associated with local opening of the double helix, a phenomenon that can be triggered *in vitro* by varying temperature, pH or salt concentration [6]. The melting transition of DNA from one double-stranded (ds) helix to two single-stranded (ss) coils has been intensively studied in the past by means of buoyant densities experiments [7], hyperchromicity spectra [8], AFM measurements [9], singlemolecule experiments [10] and fluorescence microscopy [11].

In particular, experiments [6, 7] and theories [12, 13] have shown that the “helix-coil” transition in linear or nicked DNA molecules, which do not conserve the total linking number between the two strands, is abrupt and bears the signature of a first-order-like transition. On the other hand, the same transition is much smoother for DNA segments whose linking number is topologically preserved, such as circular, covalently-closed ones [7, 14].

Understanding the physical principles underlying DNA melting in topologically constrained (tc) DNA is important since this is the relevant scenario *in vivo*. For instance, bacterial DNA is circular, while in eukaryotes DNA wraps around histones [2], and specialised proteins are able to inhibit the diffusion of torsional stress [15].

Intriguingly, and in stark contrast with the behaviour of linear, or topologically free (tf), DNA, the width of the melting transition of tcDNA is relatively insensitive to the precise nucleotide sequence [14] thereby suggesting that a universal physical mechanism, rather than biochemical details, may underlie the aforementioned broadening. While biophysical theories of tcDNA melting do exist, they do not reach a consensus as to whether the transition should weaken or disappear altogether [16–20]; more importantly, the physics underlying the broadening of the transition remains unclear.

Here we shed new light on this issue by employing a combination of complementary methods. First, we perform large-scale coarse-grained Brownian Dynamics (BD) simulations of 1000 base-pairs (bp) long topologically free and constrained double-stranded DNA molecules undergoing melting [21]. These unprecedentedly large-scale simulations at single bp resolution predict topology-dependent melting curves in quantitative agreement with experiments. Second, we propose and study a phenomenological Landau mean field theory which couples a critical “denaturation” field (*ϕ*) with a non-critical “supercoiling” one (*σ*). This approach captures the interplay between local DNA melting and topological constraints, and predicts the emergence of phase coexistence within a wide temperature range, in line with our simulations. This provides a generic mechanism to explain the observed experimental broadening of the melting transition in tcDNA. We also derive dynamical equations for the fields *ϕ* and *σ* and discuss the topology-dependent exponents describing the coarsening of denaturation bubbles during DNA melting.

## Melting Curves and Phase Diagram

We first investigate the melting behaviour of DNA by means of BD simulations of the model proposed in [21]. The dsDNA is made up by two single-stranded chains of “patchy-beads” connected by permanent FENE bonds. Every patch-bead complex represents one nucleotide and complementary strands are paired by bonds connecting patches. We model these bonds as breakable harmonic springs, which mimic hydrogen (H) bonds between nucleotides (see SM for further details). For simplicity, we consider a homopolynucleotide (no sequence dependence) [14].

As anticipated, the double-helical structure can be opened up *in vitro* either by increasing the temperature (*T*), or by increasing pH or salt concentration: both methods effectively reduce the strength of the H bonds, *ϵ*_HB_, in between nucleotides. The simulations reported in this Letter emulate the latter route: starting from an equilibrated dsDNA molecule, we perform a sudden quench of *ϵ*_HB_, and record the time evolution of the system until a new steady state is reached (see SM Fig. S2).

An observable that directly compares with experiments is the fraction of denatured base-pairs (bp), *ϑ*. The plot of the equilibrium value 〈*ϑ*〉 as a function of temperature or bond strength can be identified with the melting curve for DNA. Typical profiles obtained from experiments [7] and BD simulations performed in this work, are shown in Fig. 1(A-B): the qualitative agreement is remarkable.

**Figure 1.**
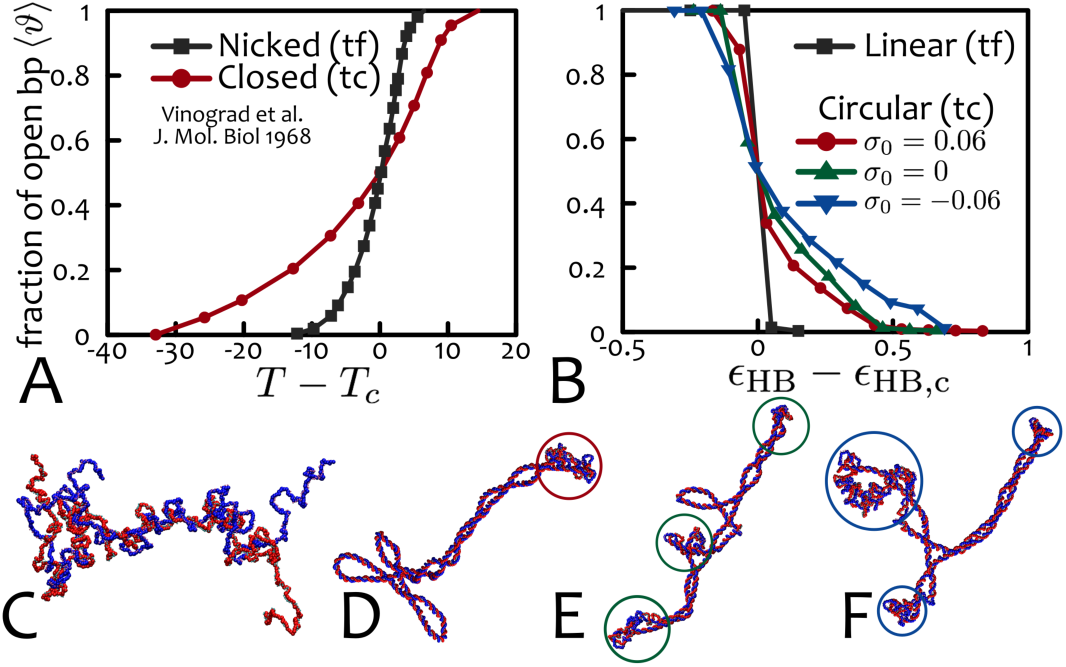
Melting curves. (**A**) shows the melting curves (fraction of denatured bp, 〈*ϑ*〉) for nicked and intact polyoma DNA as a function of *T* − *T_c_* (data from [7]). (**B**) shows the melting curves obtained in the present work via BD simulations of tf and tcDNA molecules, with length 1000 bp and different levels of supercoiling, as a function of the (shifted) effective hydrogen-bond strength *ϵ*_HB_ − *ϵ*_HB,c_ (averaged over 5 replicas and 10^6^ BD timesteps). In both experiments (**A**) and simulations (**B**), the transition appears smoother for tcDNA and the relative broadening Δ*t*|_tc_ / Δ*t*|_tf_ ∼ 3 is in quantitative agreement. The critical bond energies for which half of the base-pairs melt are *ϵ*_HB,c_/*k_B_T* = 1.35 for linear DNA and 0.309, 0.238, 0.168 for *σ*_0_ = −0.06, 0 and 0.06, respectively. From these values one can readily notice that the critical bond energy decreases (linearly) with supercoiling. (**C**)-(**F**) show snapshots of typical configurations for *ϵ*_HB_ = 0.3 *k_B_T* for tf (linear) and tcDNA with *σ*_0_ = 0.06, 0, −0.06, respectively. Stably denatured bubbles localise at regions of high curvature (tips of plectonemes [22], highlighted by circles). In (**C**) the linear DNA molecule is in a fully denatured state.

The sharpness of the melting transition can be quantified in terms of the maximum value attained by the differential melting curve as Δ*t* = |*d*〈*ϑ*〉/*dt*|^−1^, where *t* can either be temperature, *T*, or effective hydrogen bond strength, *ϵ*_HB_, depending on the denaturation protocol. Quantitatively, Figures 1(A)-(B) show that experiments and simulations agree in predicting melting curves for tcDNA about three times broader than for tfDNA, i.e. Δ*t*|_tc_ / Δ*t*|_tf_ ≃ 3.

From these observations, it is clear that the melting behaviour of DNA is affected by global topology. On the other hand, melting occurs through local opening of the double-helical structure. The challenge faced by a theory aiming to understand the “helix-coil” transition in tcDNA is therefore to capture local effects due to the global topological invariance. To this end, it is useful to define an effective local supercoiling field *σ*(*x*,*t*) ≡ (*Lk*(*x*,*t*) − *Lk*_0_) /*Lk*_0_, where *Lk*_0_ is the linking number between the two strands in the relaxed B-form state, i.e. 1 every 10.4 bp, and *Lk*(*x, t*) is the effective linking number at position *x* and time *t*. For a circular closed molecule of length *L*

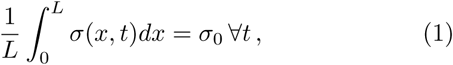
where *σ*_0_ is the initial supercoiling deficit, which can be introduced and locked into the chain by, for instance, the action of topological enzymes [2]. On the contrary, circular nicked or linear (tf) dsDNA molecules need not satisfy Eq. (1), since any deviation from the relaxed superc-oiling state can be expelled through the chain ends or the nick. In light of this, it is clear that subjecting a tcDNA to denaturation-promoting factors causes a competition between entropy and torsional stress: the former associated with the denatured coiled regions [12, 13], the latter arising in the intact helical segments [17].

Motivated by these observations, we propose the following phenomenological mean field theory for the melting of tf and tcDNA. We consider a denaturation field, *ϕ*(*x, t*), describing the state of base-pair *x* at time *t* (e.g., taking the value 0 if intact or > 0 if denatured), coupled to a conserved field, *σ*(*x, t*), tracking the local supercoiling. A Landau free energy can be constructed by noticing that: (i) the denaturation field *ϕ* should undergo a first-order phase transition when decoupled from *σ* [12], (ii) the elastic response to the torsional stress should be associated with even powers of *σ* [18] [23], and (iii) the coupling between *σ* and *ϕ* should be such that there should be an intact dsDNA phase at sufficiently low *T*, i.e. *ϕ* = 0 for any *σ*_0_ at *T* < *T*_c_.

Based on these considerations we can write an effective free energy density as:

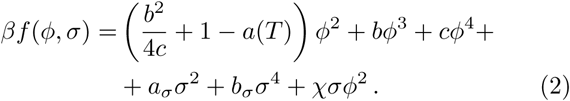

In Eq. (2), the first term is written so that the parameter *a*(*T*) ∼ *T*/*T_c_, b* < 0 and we keep the quartic term in *σ* to ensure there is a local minimum around *σ* = −1 (or *Lk*(*x, t*) = 0) when *ϕ* = *ϕ*_0_, corresponding to the fully denatured state (see SM). Finally, the coupling term χ*σϕ*^2^ models the interplay between supercoiling and local melting; it can be turned off by simply setting χ = 0 to approximate tfDNA (in this framework the torsional stress can be expelled infinitely fast). We also highlight that by setting χ = 0, Eq. (2) predicts a first-order melting transition.

The free energy density in Eq. (2) displays a minimum at *ϕ* = 0 = *ϕ*_ds_ (helical state), and can develop a competing one at *ϕ* = *ϕ*_ss_ > 1 (coiled state) which in general depends on *σ*, χ and *a* (see SM). The free energy density of the two becomes equal at the critical temperature *a* = *a_c_*, which reduces to *a_c_*(*σ*) = 1 + χ*σ*, by fixing *c* = −*b*/2 (see SM). This relation states that more negatively supercoiled molecules denature at lower temperatures, as in experiments [24], whereas for tfDNA (χ = 0) the critical temperature is independent on supercoiling.

To obtain the phase diagram of the system in the space (*a, σ*_0_) we focus on dsDNA molecules with fixed, and initially uniform, value of supercoiling *σ*_0_, at fixed temperature *a*. For such conditions, the system attains its free-energy minimising state for a value of *ϕ* = *ϕ*_0_ (χ, *σ*_0_, *a*) [25]. The uniform solution (*σ*_0_, *ϕ*_0_) is linearly unstable if it lies within the spinodal region (in Fig. 2 shaded in grey), i.e. where *∂*^2^*f*(*ϕ*_0_, *σ*)/*∂σ*^2^ ≤ 0 [26]. In Figure 2 we fix for concreteness *b* = −4, *a_σ_* = 1, *b*_σ_ = 1/2, χ = 2 (different parameter choices lead to similar diagrams provided *b* remains negative [27]).

**Figure 2.**
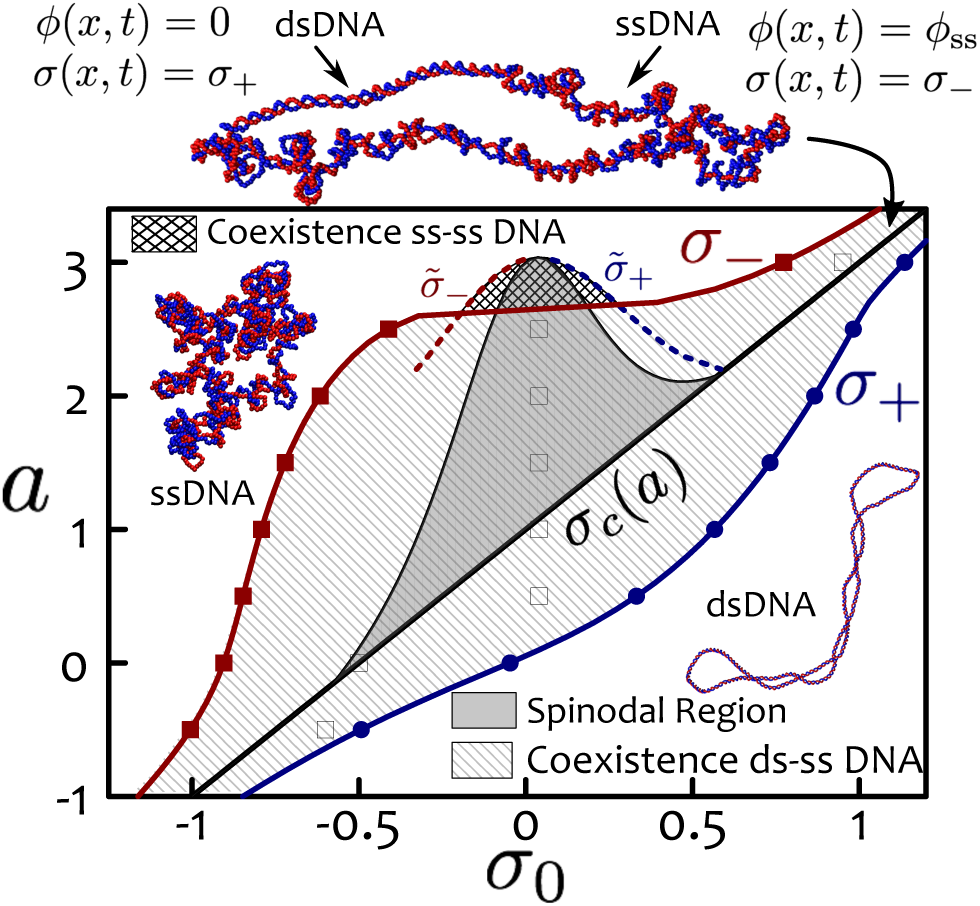
Phase Diagram. The thick solid line represents the hidden first order transition line *σ_c_*(*a*). The line-shadowed area highlights the region of absolute instability of the uniform phase; the spinodal region is coloured in grey. Binodal lines are denoted as *σ*_−_ and *σ*_+_. Cross-shadowed area highlights the region of coexistence of two denatured (ss) phases. Filled symbols denote the values obtained from numerical integration of Eq. (7), with initial *σ*_0_ as indicated by the empty squares. Snapshots of ssDNA, dsDNA and ds-ss DNA coexistence observed in BD simulations are also shown.

A system with unstable uniform solution separates into two phases with low (*σ*_−_) and high (*σ*_+_) supercoiling levels, as this lowers the overall free energy. The values of *σ*_−_ (*a*) and *σ*_+_(*a*) are the coexistence curves, or binodals, which are found by imposing that both chemical potential *μ*(*s*) ≡ *∂f*(*ϕ*_0_, *σ*)/∂*σ*|*_s_* and pressure ∏(*s*) = *f*(*ϕ*_0_, *s*) *− μ*(*s*)*s* must be equal in the two phases [25]. This translates into solving a system of two equations with two unknowns,

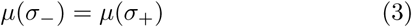

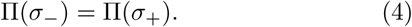

By noticing that *σ*_0_ needs to satisfy Eq. (1) for tcDNA, it is straightforward to find the fractions of the system in the high and low supercoiling phases as *f*_+_ = (*σ*_0_ − *σ*_−_)/(*σ*_+_ − *σ*_−_) and *f*_−_ = (*σ*_+_ − *σ*_0_)/(*σ*_+_ − *σ*_−_), respectively.

The phase diagram in the (*a, σ_0_*) space is reported in Figure 2, where we show that the coexistence lines *σ*_−_(*a*) and *σ*_+_(*a*) wrap around the critical first-order transition line *σ_c_*(*a*) = (*a* − 1)/χ which therefore becomes “hidden” [25]. In light of this we argue that the smoother transition observed for tcDNA [7, 14] can be understood as a consequence of the emergence of a coexistence region in the phase space which blurs the underlying first-order transition. This argument also explains the “early melting” of closed circular DNA [7], which can be understood as the entry into the coexistence region from low temperatures.

Intriguingly, our phase diagram includes a region (cross-shadowed in Fig. 2) where the system displays stable coexistence of two open phases (i.e. *ϕ* = *ϕ*_ss_ in both sub-systems) with supercoiling levels 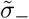 and 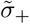.

## Dynamical Scaling

The dynamics of the non-conserved order parameter, *ϕ*, and the conserved one, *σ*, can be found following the Glauber and Cahn-Hilliard prescriptions, respectively. Consequently, the system can be described by the following “model C” equations [26]

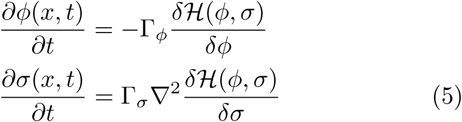
where Γ*_σ,ϕ_* are relaxation constants,

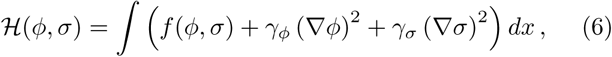
and *γ*_*ϕ*,*σ*_ determine the effective surface tension of bubbles and supercoiling domains, respectively. From Eqs. (2) and (5) one can write

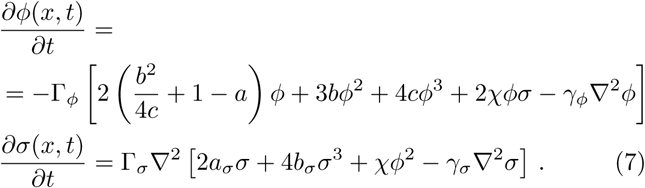

We numerically solve this set of partial differential equations (PDE) on a 1D lattice of size *L* for fixed *a* and *σ*_0_ (see SM for details) and compare the evolution of denaturation bubbles with the one observed in BD simulations. Note this set of equations disregards thermal noise, hence it is in practice a mean field theory.

In Figure 3 we show “kymographs” from BD simulations, capturing the state of each base-pair (either intact or denatured) against time for tf and tcDNA. As one can notice, after the energy quench at *t* = 0, the linear (tfDNA) molecule starts to denature from the ends and eventually fully melts. On the other hand, in the closed circular (tcDNA) molecule, bubbles pop up randomly over the whole contour length, and the steady state entails a stable fraction, 0 < *ϑ* < 1, of denatured bp (see also SM, Fig. S2).

**Figure 3.**
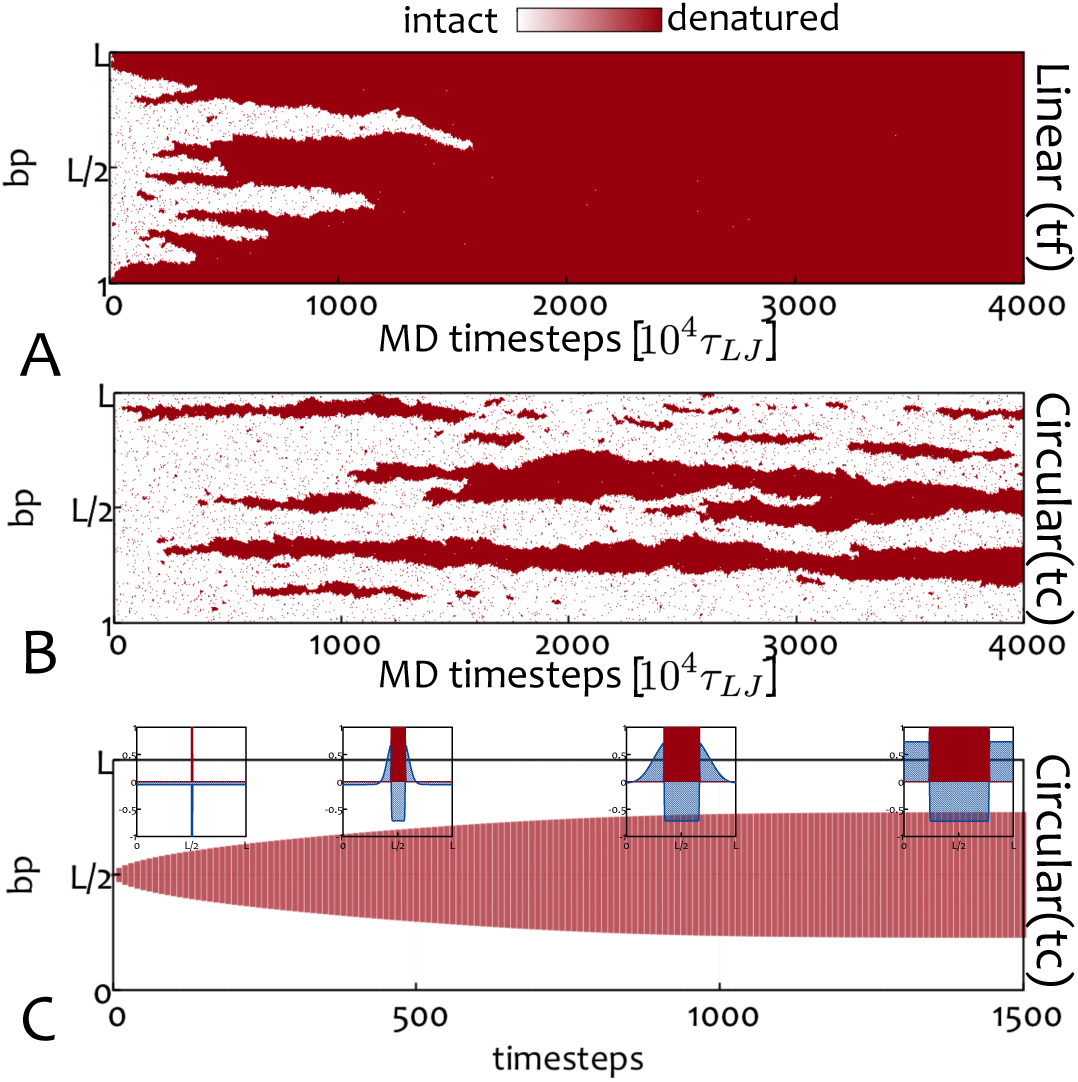
Kymographs. (**A-B**) report results from BD simulations. At time *t* = 0, the H bond strength is quenched to *ϵ*_HB_ = 0.3*k_B_T* and the local state of the chain (red for denatured and white for intact) is recorded as a “kymograph”. (**A**) and (**B**) show the case of a tf and tcDNA (*σ*_0_ = 0), respectively. (**C**) shows the kymograph of the system during integration of Eqs. (7) starting from a small bubble (see SM). Insets show instantaneous profiles of denaturation field (red) and supercoiling field (blue).

We observe a similar behaviour when the fields *ϕ* and *σ* are evolved via Eqs. (7), starting from a single small bubble at temperature *a* within the coexistence region (see Fig. **3**D and SM). While the bubble grows, the supercoiling field is forced outside the denatured regions and accumulates in the ds segments. The increasing positive supercoiling in the helical domains slows down and finally arrests denaturation, resulting in phase coexistence in steady state, between a denatured phase with *σ* = *σ*_−_ and *ϕ* = *ϕ*_ss_ > 1, and an intact phase with *σ* = *σ*_+_ and *ϕ* = *ϕ*_ds_ = 0.

The growth, or coarsening, of a denaturation bubble, *l*, can be quantified within our mean field theory and BD simulations: in the former case by numerical integration of Eq. (7), in the latter by measuring the size of the largest bubble over time and averaging over independent realisations. As shown in Fig. 4(A-C) we find that in both mean field and BD simulations,

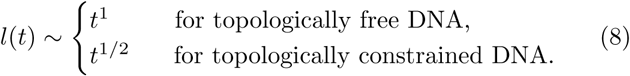

**Figure 4.**
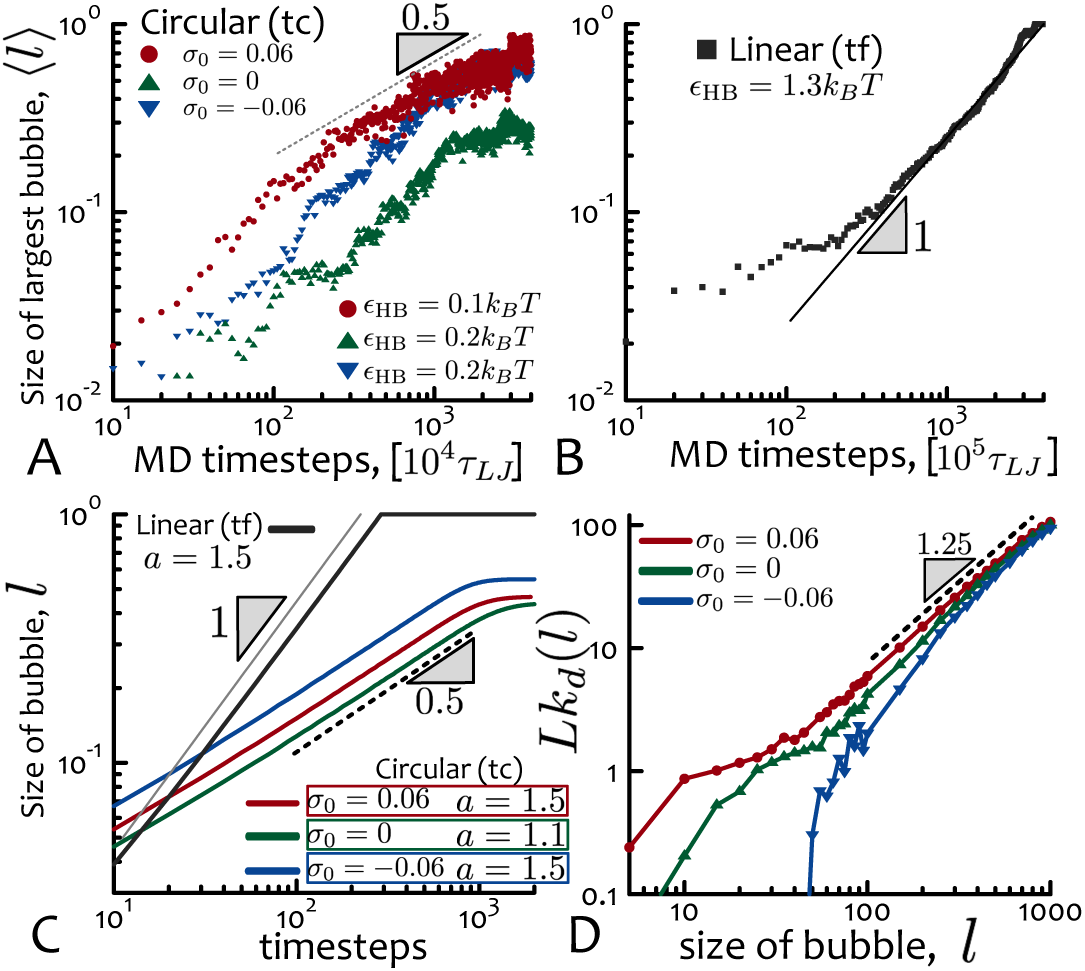
Dynamical Scaling. (**A-B**) show results from BD simulations. The size of the largest denatured bubble 〈*l*〉 (averaged over 5 replicas) is plotted against time from the moment of the quench. (**A**) shows tcDNA while (**B**) refers to tfDNA. (**C**) shows the size of a single growing bubble, *l*, within our mean field model, Eqs. (7). PDE and BD simulations show similar behaviours, which suggest a universal dynamical scaling with topology-dependent exponent (α = 1 for χ = 0 and *α* = 1/2 for χ > 0). (**D**) shows the linking number, *Lk_d_*, stored inside a denatured bubble of fixed size *l* computed from BD simulations (see text and SM for details).

In other words, we find that the exponent *α* governing the *local* growth of a denaturation bubble depends on the *global* topology of the molecule.

We propose the following argument to explain the values of *α*. For tfDNA (e.g., nicked or linear), we can assume that the supercoiling field relaxes quickly, and gets expelled outside, without affecting the dynamics of the denaturation field. In this case, the free energy can be approximated as *f* ≃ (*ϵ*_HB_ − *T*Δ*S*)*l*, so that there is a constant increase in entropy per each denatured bp when *T* > *T_c_* = *ϵ*_HB_/Δ*S*. This implies that [26] *ψdl*/*dt* ≃ *df*/*dl* ~ *const*, with *ψ* an effective constant friction; as a result we obtain *l*(*t*) ∼ *t*.

On the other hand, the value of *α* = 1/2 observed for tcDNA (e.g., circular non-nicked plasmids) can be understood by quantifying the slowing down of denaturation due to the accumulation of a “wave” of supercoiling, raked up on either side of the growing bubble. We argue that the flux of *ϕ* through a base pair at the bubble/helix interface is *J_ϕ_* ∼ *ϕ_ss_dl*/*dt*. At the same time, the flux of *σ* can be obtained by noticing that the “wave” can be approximated by a triangle with constant height *h* = *σ*_+_ – *σ*_0_ and base *b* ∼ *l*(*t*) (see SM Fig. S10 and Fig. 3(C)). This is because the total supercoiling enclosed by the wave must be proportional to the one expelled from within the denatured bubble, which is *∼* |*σ*_−_ − *σ*_0_| *l*(*t*). One can therefore write that *J_σ_* = −Γ*_σ_∂_x_σ* ≃ Γ_*σ*_*h*/*l*, for the supercoiling flux in, say, the forward direction. At equilibrium, the two fluxes must balance, i.e. *J_ϕ_* ∼ *J_σ_*, and therefore *ϕ*_ss_*dl*/*dt* ∼ Γ*_σ_h*/*l*, or *l*(*t*)∼*t*^1/2^. Finally, we highlight that this argument depends on the slowing down over time of 1D supercoiling fluxes, hence is qualitatively distinct from the reason why *α* =1/2 in dimensions *d* ≥ 2 in (non-conserved) model A [26].

## Linking within Denaturation Bubbles

As a final result, we perform BD simulations to characterise the topology of a denaturation bubble through the linking number that can be stored inside it (Fig. **4**(D)). An idealised bubble is identified by Lk = 0 (and *σ* = −1) [17], whereas our BD simulations show that a denatured region of *fixed* size *l* (imposed by selectively breaking *only l* consecutive bonds along an intact dsDNA molecule) has a non-zero linking number *Lk_d_* (see Fig. **4**(D)) [28].

We found that for small *l, Lk_d_* displays a remarkable signature of global topology (through the value of *σ*_0_); instead, the scaling behaviour at large *l* appears to follow *Lk_d_* ∼ *l*^1.25^ irrespectively of *σ*_0_, until it reaches *Lk*_0_. The finding that a denaturation bubble displays a non-trivial and *l*-dependent linking number suggests that idealised (*Lk_d_* = 0) bubbles may not always be reflecting realistic behaviour. Further, it may be of relevance for processes such as DNA replication, as it suggests that supercoiling or torsional stress may be able to diffuse past branching points such as replication forks [29].

## Conclusions

In summary, we have studied the melting behaviour of topologically constrained DNA through a combination of large-scale BD simulations and mean field theory. A key result is that the phase diagram for tcDNA melting generally involves a phase coexistence region between a denatured and an intact phases, pre-empting a first-order denaturation transition as in tfDNA. This finding provides a theoretical framework to explain the long-standing experimental observation that the denaturation transition in circular, and not nicked, supercoiled plasmids is seemingly less cooperative (smoother) than for linear, or nicked, DNA [7, 14].

We have further studied, for the first time, the coarsening dynamics of denaturation bubbles in tcDNA, and found a remarkable agreement between BD simulations and mean field theory, both reproducing similar topology-dependent scaling exponents that can be understood within our theoretical model. It would be of interest to investigate such dynamics experimentally in the future.

DMa and DMi acknowledge ERC for funding (Consolidator Grant THREEDCELLPHYSICS, Ref. 648050). YAGF acknowledges support form CONACyT PhD grant 384582.

## References

[1] A. Bates and A. Maxwell, DNA topology (Oxford University Press, 2005).

[2] B. Alberts, A. Johnson, J. Lewis, D. Morgan, and M. Raff, Molecular Biology of the Cell (Taylor & Francis, 2014) p. 1464.

[3] K. Rippe, P. H. von Hippel, and J. Langowski, Trends Biochem. Sci. 20, 500 (1995).

[4] L. F. Liu and J. C. Wang, Proc. Natl. Acad. Sci. USA 84, 7024 (1987).

[5] C. A. Brackley, J. Johnson, A. Bentivoglio, S. Corless, N. Gilbert, G. Gonnella, and D. Marenduzzo, Phys. Rev. Lett. 117, 018101 (2016).

[6] R. M. Wartell and A. S. Benight, Phys. Rep. 126, 67 (1985).

[7] J. Vinograd, J. Lebowitz, and R. Watson, J. Mol. Biol. 33, 173 (1968).

[8] D. E. Jensen and P. H. von Hippel, J. Biol. Chem. 251, 7198 (1976).

[9] J.-H. Jeon, J. Adamcik, G. Dietler, and R. Metzler, Phys. Rev. Lett. 105, 208101 (2010).

[10] R. Vlijm, J. v. d. Torre, and C. Dekker, PLoS One 10, e0141576 (2015).

[11] G. Altan-Bonnet, A. Libchaber, and O. Krichevsky, Phys. Rev. Lett. 90, 138101 (2003).

[12] D. Poland and H. A. Scheraga, J. Chem. Phys. 45, 1464 (1966).

[13] Y. Kafri, D. Mukamel, and L. Peliti, Phys. Rev. Lett. 85, 4988 (2000).

[14] A. V. Gagua, B. N. Belintsev, and L. Lyubchenko Yu, Nature 294, 662 (1981).

[15] C. Naughton, N. Avlonitis, S. Corless, J. G. Prendergast, I. K. Mati, P. P. Eijk, S. L. Cockroft, M. Bradley, B. Ylstra, and N. Gilbert, Nat. Struct. Mol. Biol. 20, 387 (2013).

[16] J. Rudnick and R. Bruinsma, Phys. Rev. E 65, 2 (2002).

[17] A. Kabakçiolu, E. Orlandini, and D. Mukamel, Phys. Rev. E 80, 1 (2009), 0811.3229.

[18] C. J. Benham, Proc. Natl. Acad. Sci. USA 76, 3870 (1979).

[19] S. Sen and R. Majumdar, Biopolymers 27, 1479 (1988).

[20] W. Bauer and C. Benham, J. Mol. Biol. 234, 1184 (1993).

[21] Y. A. G. Fosado, D. Michieletto, J. Allan, C. A. Brackley, O. Henrich, and D. Marenduzzo, Soft Matter 12, 9458 (2016).

[22] C. Matek, T. E. Ouldridge, J. P. K. Doye, and A. a. Louis, Sci. Rep. 5, 7655 (2014).

[23] This is a good approximation for small σ.

[24] V. Víglaský, M. Antalík, J. Adamcík, and D. Podhradský, Nucleic acids research 28, E51 (2000).

[25] A. Matsuyama, R. M. L. Evans, and M. E. Cates, Eur. Phys. J. E 87, 79 (2002).

[26] P. Chaikin and T. Lubensky, Principles of Condensed Matter Physics (Cambridge University Press, 2000).

[27] Given these values of *b, aσ, bσ*, we further require 1.72 ≤ *χ* ≤ 2.67 to obtain a spinodal region.

[28] In practice, *Lk_d_* was obtained by removing the linking number of the chain *outside* the denatured region *Lk_out_* from the initially set *Lk*, see SM for details.

[29] L. Postow, N. J. Crisona, B. J. Peter, C. D. Hardy, and N. R. Cozzarelli, Proc. Natl. Acad. Sci. USA 98, 8219 (2001).

